# Identifying the abundant and active microorganisms common to full-scale anaerobic digesters

**DOI:** 10.1101/104620

**Authors:** Rasmus H. Kirkegaard, Simon J. McIlroy, Jannie M. Kristensen, Marta Nierychlo, Søren M. Karst, Morten S. Dueholm, Mads Albertsen, Per H. Nielsen

## Abstract

Anaerobic digestion is widely applied to treat organic waste at wastewater treatment plants. Characterisation of the underlying microbiology represents a source of information to develop strategies for improved operation. To this end, we investigated the microbial community composition of thirty-two full-scale digesters over a six-year period using 16S rRNA gene amplicon sequencing. Sampling of the sludge fed into these systems revealed that several of the most abundant populations were likely inactive and immigrating with the influent. This observation indicates that a failure to consider immigration will interfere with correlation analysis and give an inaccurate picture of the active microbial community. Furthermore, several abundant OTUs could not be classified to genus level with commonly applied taxonomies, making inference of their function unreliable. As such, the existing MiDAS taxonomy was updated to include these abundant phylotypes. The communities of individual plants surveyed were remarkably similar – with only 300 OTUs representing 80% of the total reads across all plants, and 15% of these identified as likely inactive immigrating microbes. By identifying the abundant and active taxa in anaerobic digestion, this study paves the way for targeted characterisation of the process important organisms towards an in-depth understanding of the microbial ecology of these biotechnologically important systems.

## Introduction

Biogas production from the anaerobic digestion of organic waste is increasingly being implemented as an alternative renewable energy source. This change is driven by the need for clean energy as well as improved economy of wastewater treatment plants by making them into net energy producers^1^. Methane gas production from organics is mediated by the tightly coupled synergistic activities of complex microbial communities and is essentially covered by four sequential stages: hydrolysis, fermentation, acetogenesis and methanogenesis. The anaerobic digestion process is generally robust, but occasionally reactors experience operational problems such as foaming events and periods of low efficiency or failure^2–4^. A better understanding of the underlying microbiology will facilitate optimisation of the biological processes, and consequently the microbiology has been widely studied using various approaches with both lab and full-scale systems^5–7^.

Understanding the ecology of anaerobic digesters, and how it relates to system function, first requires the identification of the active and abundant microorganisms and subsequent linkage of their identity to their functional roles^8^. Several 16S rRNA gene amplicon based studies have shown that there appears to be a set of abundant microorganisms, common to similarly operated anaerobic digesters, that are stably present over time^6, 7, 9, 10^. This is also known for other biological processes, such as wastewater treatment plants^11^ and the human digestive system^12^. Furthermore, other studies have revealed process temperature, substrate composition, and ammonia concentrations as important factors in the shaping of the microbial community composition. However, in anaerobic digesters a large part of the observable microbial community might originate from dead or inactive cells arriving with the influent biomass from which DNA still persists. Hence, the observed microbial community dynamics will not truly reflect the changes in process performance or stability. This can lead to spurious correlations and false conclusions^11^. In an attempt to mitigate the problem of DNA from inactive cells influencing microbial analysis, molecular techniques have been developed to remove or bind the extracellular DNA prior to cell lysis^13, 14^. However, the complex matrix of anaerobic digester sludge samples will likely lead to problems with unwanted chemical reactions and limited penetration of the light used in the process^14^. Hence, an alternative approach is to monitor the microbial composition in the influent to identify organisms whose abundance is likely maintained by immigration^11, 15–17^.

Associating phylogeny with function is essential for understanding the ecology of these systems. However, a substantial proportion (67%^9^ to 73%^7^) of sequences obtained in previous 16S rRNA gene amplicon surveys of anaerobic digesters were not classified to the genus level with the commonly applied taxonomies: such as SILVA, RDP and Greengenes^18–20^. Furthermore, biases associated with DNA extraction, primer coverage and differences in the taxonomy applied for classification^21, 22^, greatly hampers cross-study comparisons. Hence, only by using well-defined standard methods and the same curated database for taxonomic classification across the field, it is possible to make meaningful cross-study comparisons and robust biological conclusions^23^. Standardisation has been established for activated sludge from wastewater treatment plants with the MiDAS protocols^21^ and the curated MiDAS taxonomy^22^, but is currently lacking for anaerobic digestion.

The aim of this study was to identify the abundant and active organisms in full-scale anaerobic digester systems, fed waste activated sludge, using 16S rRNA gene amplicon sequencing. The survey included 32 Danish full-scale reactors located at 20 wastewater treatment plants over a six-year period (>300 samples), including both mesophilic and thermophilic reactors, and represents the most comprehensive study of full-scale systems to date. Comparison of abundances in the digester sludge, and the corresponding influent primary and surplus sludge, was used to identify immigrating populations and to provide an assessment of the activity of populations in the anaerobic digesters. Furthermore, having identified the abundant populations present in the anaerobic digesters, we have performed a manual curation of the SILVA taxonomy for the most abundant OTUs, many of which were poorly classified with existing databases. By providing genus level classifications for all abundant taxa, researchers in the field will be able to link the identity with the accumulated knowledge regarding their population dynamics and ecophysiology.

## Results

### Characteristic of the sampled anaerobic digesters

More than 300 samples were collected from 32 full-scale digesters at 20 wastewater treatment plants in Denmark over a period of 6 years (2011-2016). The sampled reactors represent mesophilic (~37°C) and thermophilic (~55°C) processes running mainly on primary sludge and surplus activated sludge (approx. 50:50% in relative amount). The reactors have reported ammonium levels in the range of 500-3000 mg/L, VFA concentrations of 0.5-20 mmol/L, alkalinity levels of 0.01-0.5 mmol/L, pH of 7.1-8.5, reactor volumes of 1300-6000 m^3^, and sludge retention times of 10-55 days. The plants in Fredericia and Næstved have mesophilic reactors with a thermal hydrolysis pre-treatment (THP), the type of pre-treatment in both cases are CambiTHP^TM^ installations.

### Community structure: Archaea

The archaea were targeted with archaea-specific primers amplifying the V3-5 regions of the 16S rRNA gene. The resulting quality filtered sequencing data were subsampled to 10 000 reads per sample, giving more than 3 million reads in total. There were 169 OTUs, spanning 8 phyla, which constituted at least 0.1% in a single sample. Principal component analysis revealed that the thermophilic and mesophilic reactors formed very distinct archaeal communities (**Fig. 1A**). Euryarchaeota was by far the most dominant archaeal phylum making up 93-100% of the archaeal reads in each sample (**Fig. 2A**). The acetoclastic methanogenic genus *Methanosaeta* dominated the mesophilic reactors (60-80% of the reads), followed by a variety of hydrogenoclastic methanogenic genera such as *Methanolinea*, *Methanospirillum*, *Methanobrevibacter* as well as the WCHA1-57, which was recently renamed *Candidatus* Methanofastidiosa^24^ (**Fig. 2B**). The mesophilic reactors with thermal hydrolysis pre-treatment were also dominated by *Methanosaeta* (83-87 %). The underlying OTUs for the most abundant genera were the same for the different plants (**Fig. S1**). For the genera *Methanosaeta*, there was 1 dominant OTU (25-33% relative abundance) and 6 additional OTUs in abundances of each 2-15% in all mesophilic reactors, including those with THP, indicating a substantial diversity within the genera.

**Figure 1.**
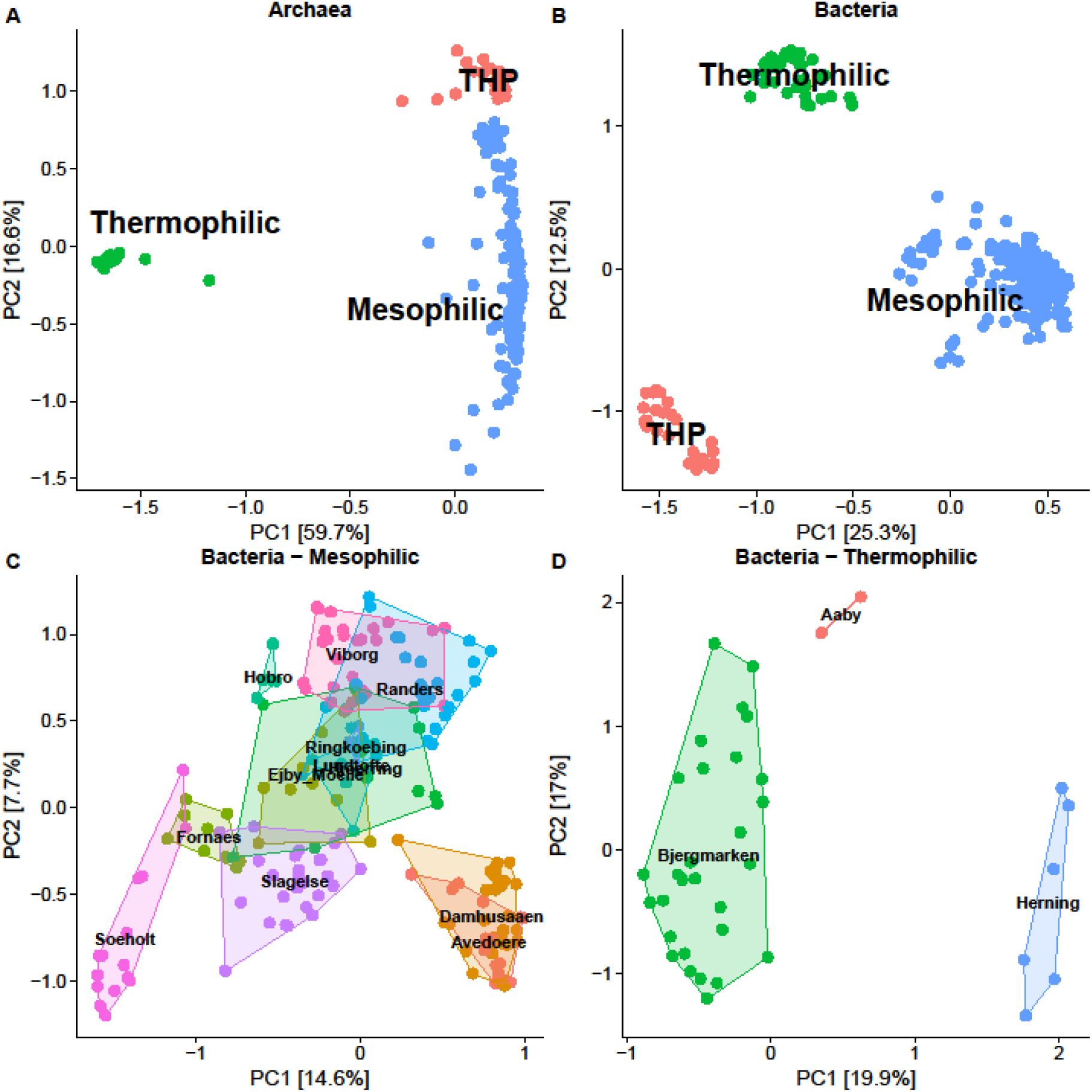
Principal component analysis of the microbial communities in ADs, highlighting samples by process type information (• mesophilic, • thermophilic, • mesophilic with thermal hydrolysis pretreatment (THP)). A) the separation of archaeal communities coloured by process type, B) the separation of bacterial communities coloured by process type, C) The bacterial communities of mesophilic plants coloured and labelled by plant location D) The bacterial communities of thermophilic plants coloured and labelled by plant location.

**Figure 2.**
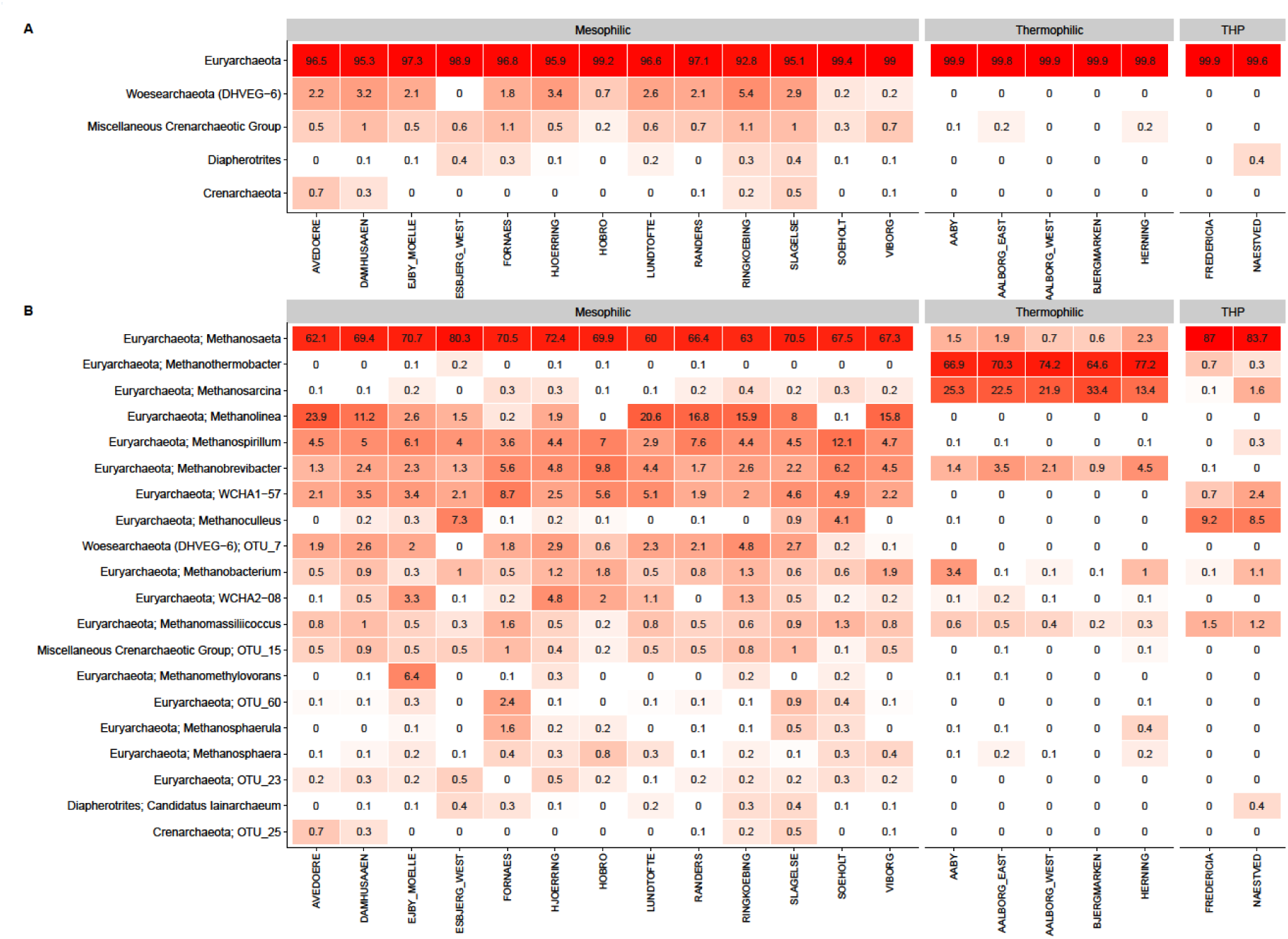
A) Heatmap of the 5 most abundant archaeal phyla. B) Heatmap of the 20 most abundant archaeal genera in the anaerobic digesters. When no genus level classification is available the OTU number is given. The phylum level classification is shown for all genera. Data based on 32 AD reactors (1-4 per plant) analysed 2-23 times. The mean abundance is shown for each plant. The taxa are sorted by mean abundance across the plants at the respective phylogenetic level (phylum, genus).

The thermophilic reactors were dominated by the hydrogenoclastic methanogenic genus *Methanothermobacter* (64-77% of the reads), followed by the more versatile *Methanosarcina* (13-33% of the reads). The latter is known to perform both acetoclastic and hydrogenoclastic methanogenesis. *Methanobrevibacter* was the third most common methanogen and along with *Methanosaeta* the only abundant archaeon shared with the mesophilic reactors. However, it was not found in mesophilic reactors with thermal hydrolysis pre-treatment. The underlying OTUs for the two abundant genera were the same for the different plants (**Fig. S1**). For *Methanothermobacter*, there was 1 dominant OTU (37-48% relative abundance) and 2 less abundant OTUs (6-20%). For *Methanosarcina*, there was 1 dominant OTU (10-25%) and 1 less abundant OTU (3-6%). The archaeal community of the thermophilic samples clearly had a lower diversity than the mesophilic samples **(Fig. 2B)**.

### Community structure: Bacteria

The bacteria were targeted with bacteria-specific primers amplifying the V1-3 regions of the 16S rRNA gene. The resulting quality filtered sequencing data were subsampled to 10 000 reads per sample giving more than 3 million reads in total. The resulting 5614 OTUs, each making up at least 0.1% of the reads in at least one sample, covered 46 phyla. Principal component analysis revealed that the thermophilic and mesophilic reactors formed very distinct bacterial communities with a separate cluster for reactors with thermal hydrolysis pre-treatment (**Fig. 1B**). Principal component analysis of the samples within the mesophilic and thermophilic clusters **(Fig. 1C & 1D**) show that the overall structure of the microbial communities overlap between some plants during the period. The dominant phyla were Firmicutes, Proteobacteria, Actinobacteria, Bacteriodetes and Chloroflexi (**Fig. 3A**). Along with the more “well-known” phyla, a few candidate phyla, such as Fermentibacteria (Hyd-24-12), Aminicenantes (OP8) and Atribacteria (OP9), were also observed. Most mesophilic reactors were dominated by the MiDAS genus T78 belonging to Chloroflexi, followed by the genera *Tetrasphaera* and *Ca.* Microthrix (**Fig. 3B**). The thermophilic reactors also had a high abundance of *Tetrasphaera* and *Ca.* Microthrix. However, the mesophilic reactors with thermal hydrolysis pre-treatment did not have a notable abundance of either of these two genera despite them being present in the surplus sludge. This suggests that these genera do not grow in conventional mesophilic digesters, but are coming in with the feed. Supporting this idea is that the underlying OTUs for the most abundant genera were the same for the different plants (**Fig. S2**). The dominant OTUs in the digesters were generally shared among the plants with similar operation (**Fig. S3**) and as few as 300 OTUs account for 80% of the reads, which is a metric sometimes defined as the “abundant core” (**Fig. S4**)^11^.

**Figure 3.**
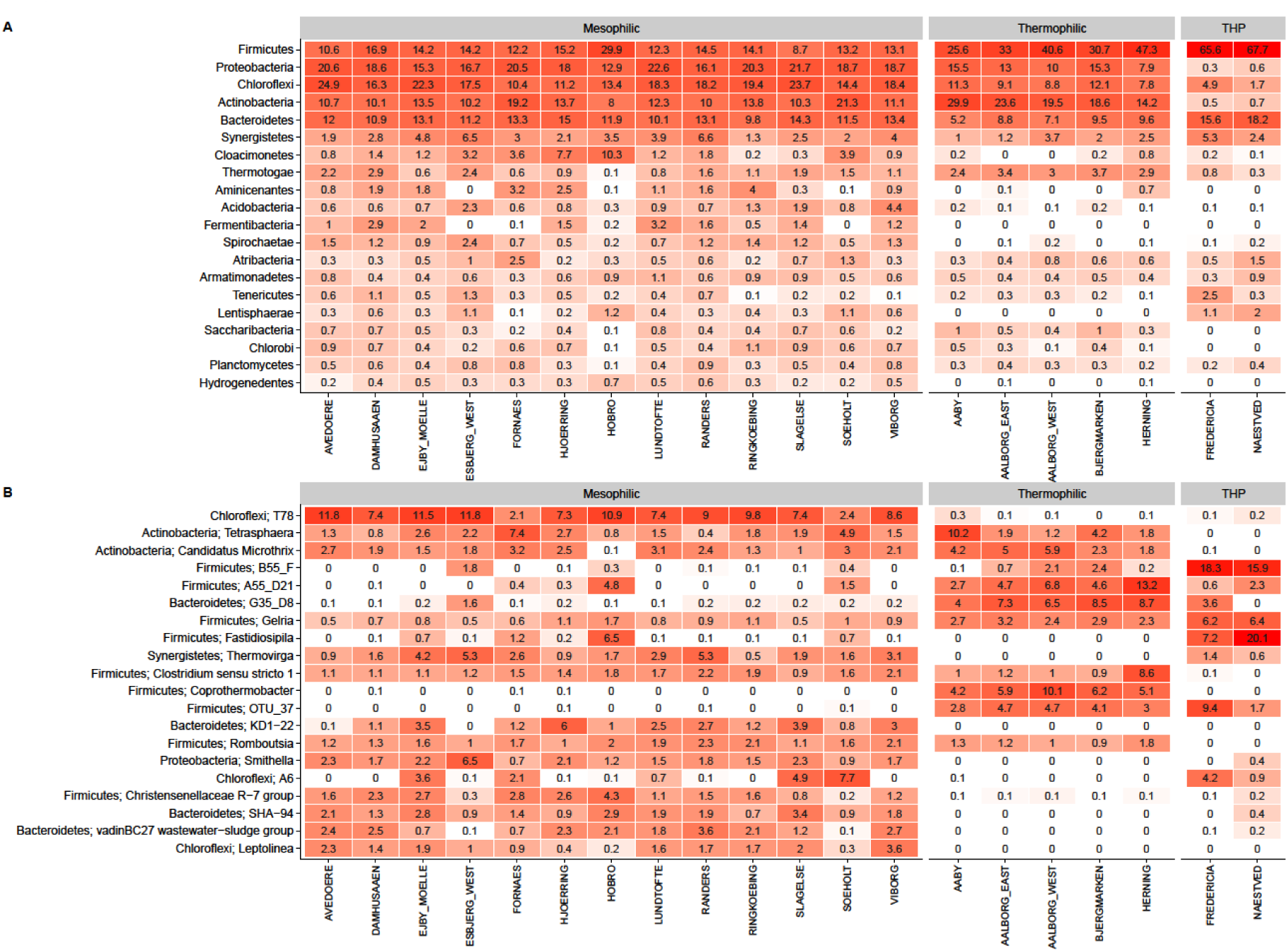
A) Heatmap of the 20 most abundant bacterial phyla. B) Heatmap of the 20 most abundant bacterial genera in the anaerobic digesters. When no genus level classification is available the OTU number is given. The phylum level classification is shown for all genera. Data based on 32 AD reactors (1-4 per plant) analysed 3-30 times. The mean abundance is shown for each plant. The taxa are sorted by mean abundance across the plants at the respective phylogenetic level (phylum, genus).

### Community composition of primary and surplus sludge

The feed for all digesters, except Fredericia, was a mixture of primary sludge settled from influent wastewater and surplus sludge harvested from the activated sludge plant, approximately in a mass ratio of 50:50. The bacterial community composition was analysed in 121 samples of primary sludge from 14 WWTPs and 137 activated sludge samples from all 24 WWTPs. The overall community structure showed clear clustering of the different sample types, separating primary sludge, surplus sludge, mesophilic, thermophilic and THP reactors (**Fig. S5**), indicating noticeably different communities. The microbial communities in the primary sludge were very similar in all samples and the most abundant genera were *Streptococcus, Arcobacter*, and *Trichococcus* (**Fig. S6**). The most abundant genera in the surplus sludge were also very similar in most plants reflecting the presence of abundant core species such as *Tetrasphaera, Ca.* Microthrix, and *Ca.* Amarilinum (**Fig. S7**).

### Survival of influent bacteria in the digesters

Some organisms were present in both of the influent streams and the digesters, whereas others were detected almost exclusively in one of the three sample types (**Fig. 4 & 6**). No overlap was found between the communities in the influent streams and in reactors with THP (**Fig. 4**). Some organisms, such as *Tetrasphaera*, *Ca.* Microthrix and *Rhodobacter*, were generally present in both the surplus sludge and the digesters, regardless of process temperature. Other organisms, such as *Arcobacter, Streptococcus*, and *Blautia*, which were the most abundant bacterial genera in the primary sludge, were hardly detected in the digesters.

**Figure 4.**
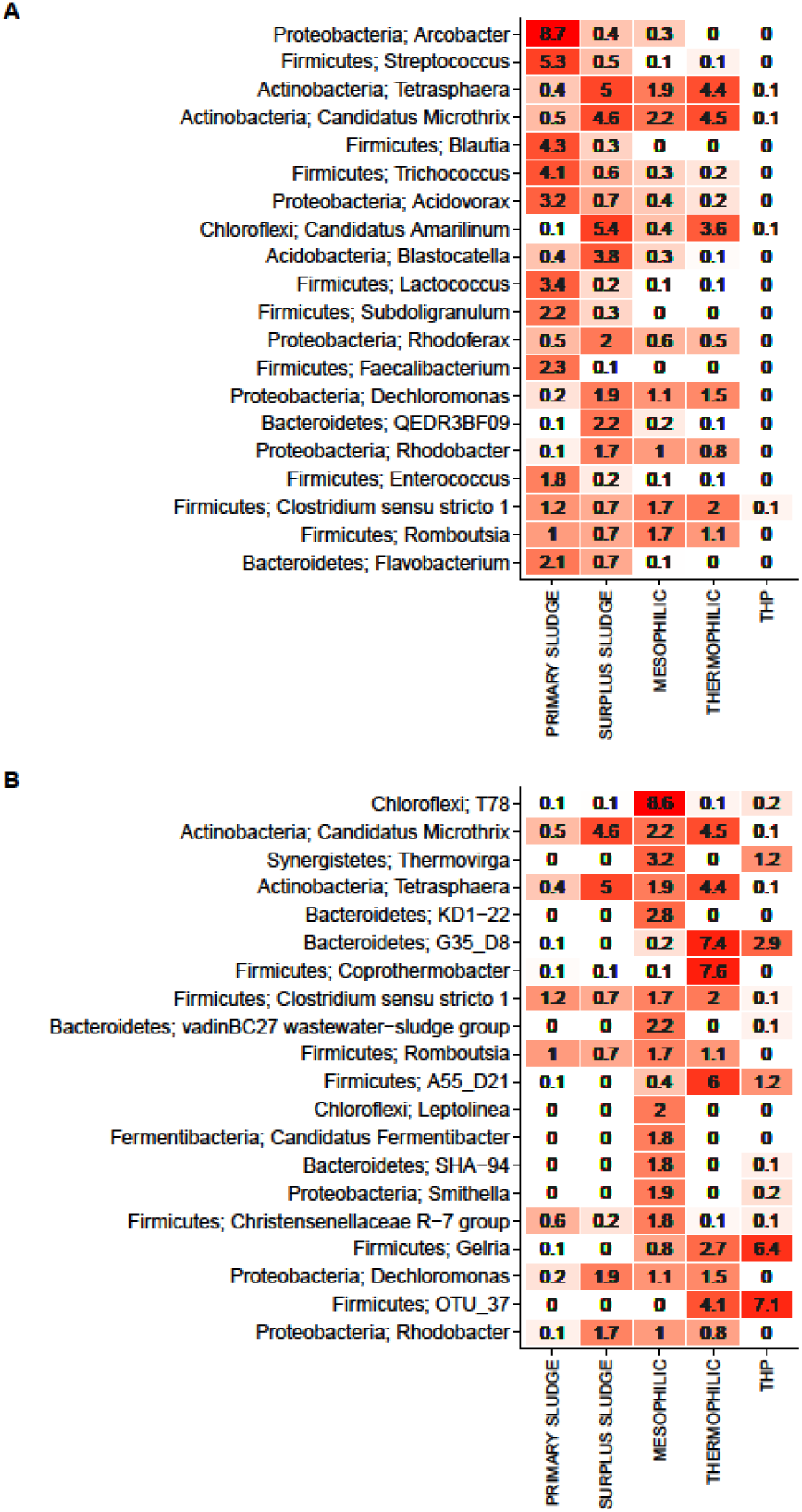
Heatmap of the 20 most abundant bacterial genera A) Taxa sorted by the mean abundance in the influent (primary and surplus sludge) B) Taxa sorted by the mean abundance in the anaerobic digesters (mesophilic, thermophilic and THP). The numbers represent mean abundances for groups with more reactors and more samples (30-279).

We tried to assess whether the immigrating organisms tended to die off, survive or grow in the digesters by calculating the ratios of their mean abundance in the three digester types compared to the mean abundance in the influent streams (**Fig. 5, Fig. 6 & Fig. S3 & Table S2**). This calculation does not give an exact measure of the growth rate of the individual species as has previously been performed based on detailed mass-balances^11,17^. However, despite some variability in the sludge retention times and in the fraction of primary and surplus sludge, etc. a clear bimodal distribution of the ratios (**Fig. 6**) was observed with a split around 10, indicating that there was a clear difference between organisms growing exclusively in anaerobic digesters and organisms that were dying off or only present because they were fed into the digester. Some of the seemingly most abundant organisms in the digesters, such as the genera *Tetrasphaera* and *Ca.* Microthrix, had ratios close to or below one (**Fig. 5**), which indicate that they were likely not growing and likely only present by being supplied with the influent streams. However, 203 of the 300 most abundant OTUs had ratios above 10, which indicate that there is a clear shift for microorganisms growing exclusively in the digesters, e.g. the genera *Ca.* Fermentibacter, *Fastidiosipila*, and *Coprothermobacter*. Among the top 100 bacterial OTUs in the digesters, 12 had ratios below 1 and 31 had ratios below 10, indicating that the immigrating bacteria had a major impact on the apparent community composition (**Fig. S3**).

**Figure 5.**
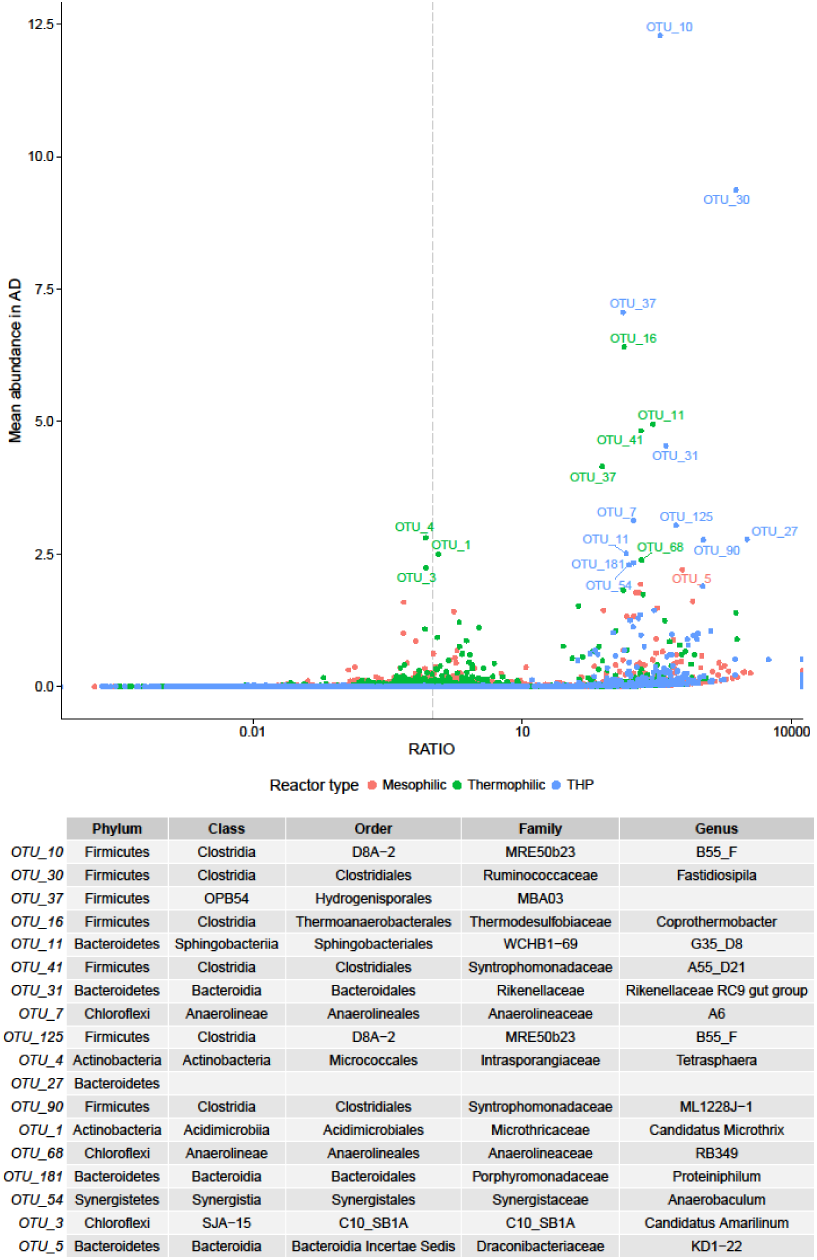
Ratio of bacterial read abundance between the digester and influent abundance versus abundance (%) in the digester for mesophilic (•), thermophilic (•), and mesophilic with thermal hydrolysis pre-treatment (• THP). OTUs with a mean read abundance above 2% are labelled with OTU numbers. Classifications of the OTUs can be found in the table below and in Table S2.

**Figure 6.**
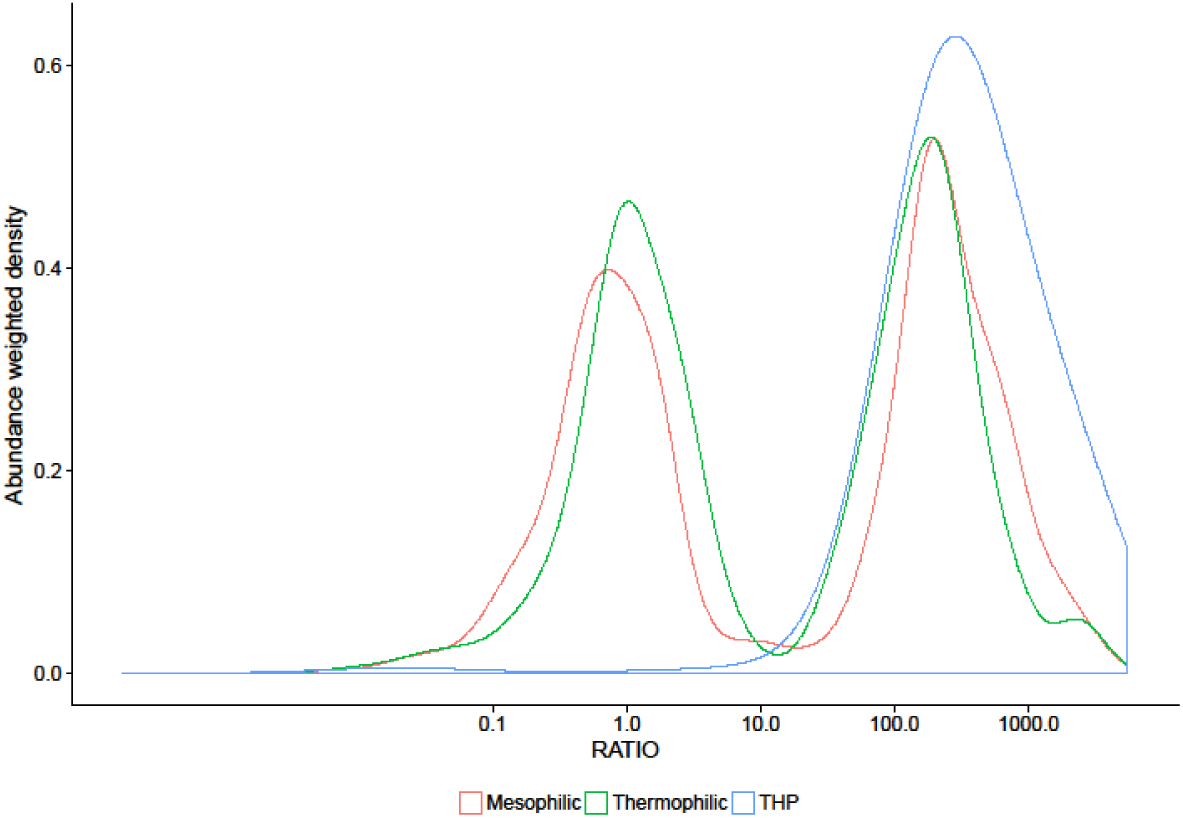
Density plot for ratios between OTU abundance in the digesters and the influent streams (primary and surplus sludge) weighted by the digester abundance.

## Discussion

In this study, the microbial communities of 32 full-scale anaerobic digesters at wastewater treatment plants and their influent streams were analysed using 16S rRNA gene amplicon sequencing, to identify the abundant and active microorganisms of these biotechnologically important systems. Principal component analysis (PCA) revealed that the bacterial communities were distinct for the thermophilic, mesophilic, and mesophilic with THP systems (**Fig. 1B**). The different communities observed in the mesophilic systems, with and without THP, may be partly attributed to the reported higher concentrations of ammonia in the latter (**Table S1**). These findings are consistent with previous studies which have also shown communities to be clearly influenced by process temperatures and ammonia concentrations^9^. The six-year survey period of the current study indicates that the digester communities at each wastewater treatment plants were relatively stable over time. Furthermore, more detailed analyses of the taxa revealed that the most abundant organisms were shared between reactors of the same process type (**Fig. 2 & 3**).

Previous studies have also observed abundant organisms that were shared among many anaerobic digester plants of similar operation^6,7,9^. This common finding indicates that efforts to characterise the process-important organisms is feasible, with less than 300 OTUs accounting for 80% of the amplicon reads across all plants in the current study. Previous attempts to identify the important genera in anaerobic digesters has been hampered by the lack of taxonomic classification of several abundant organisms with commonly applied taxonomies. As a consequence, previous studies have often focused microbial analysis on OTUs with taxonomic classification to high levels, such as phylum, order or class^6,7,9^, where the link between phylogeny and function is unreliable^25^. In this study, we have sought to address this problem by updating the MiDAS taxonomy to cover abundant genus level taxa in full-scale anaerobic digesters, along with abundant organisms previously identified in activated sludge^22^. Application of the updated taxonomy in this study gave genus level classification for 78% and 97% of all the bacterial and archaeal reads, respectively. Of the bacterial OTUs within top 300 (“abundant core”) the ones with MiDAS specific genus classification accounted for 31 % of the bacterial reads.

Importantly, a substantial presence of incoming organisms or their DNA in the community of the assessed digesters was observed in this study, indicating that some of the seemingly most abundant organisms were related to influent streams rather than actively growing. To assess the source and activity of abundant organisms, we performed the microbial analysis on the primary and surplus sludge and calculated the ratio of their abundance in these influent streams and the receiving digesters. The ratios indicate if continuous transfer into the system, and/or active growth, maintains an organism’s abundance. Fifteen percent of the 300 OTUs, which accounted for 80% of the reads, had ratios of one or below. Four of the 25 most abundant genera (**Fig. 3B**) had low relative abundance ratios. These included *Tetrasphaera*, *Ca.* Microthrix, *Clostridium sensu strictu 1*, and *Romboutsia*; which are all genera that were also seemingly shared among mesophilic and thermophilic reactors but not present in the reactors with THP. The suggestion that some of these do not belong in anaerobic digesters is also supported by what is known about their metabolism e.g. *Ca.* Microthrix is a known aerobe^26^. A similar approach also identified abundant inactive influent organisms in a single anaerobic digester treating surplus sludge^17^. Inactive organisms identified in the previous study, including *Trichococcus, Rhodobacter*, and *Thauera*, were also determined to be inactive in the current study - having ratios lower than one.

The impact of the influent on the observed community causes a multitude of problems for microbial analysis in digesters as it interferes with attempts to establish relationships between microorganisms and process performance. It is worth noting that, despite likely being inactive, the persistence of the filamentous members of the genus *Ca.* Microthrix has been linked to foaming problems in receiving anaerobic digester systems^27,28^.

The identification of inactive populations in anaerobic digesters further shortens the list of microorganisms likely important to the bulk transformations of these systems. The majority of previously characterised actively growing genera are known to be fermentative organisms; including *Coprothermobacter*^29^ and *Anaerobaculum*^30^ in thermophilic systems, and *Thermovirga*^31^*, Leptolinea*^32^, and *Ca.* Fermentibacter^33^ in mesophilic systems. *Smithella*^34^ and *Gelria*^35^ represent known acetogens. In general, apart from the influent organisms, abundant genera were generally not shared between thermophilic and mesophilic systems. The exception within the top 25 is the genus *Gelria* – which is present in both mesophilic and thermophilic reactors with a high abundance and ratio. The genus was originally isolated from a thermophilic methanogenic enrichment^36^. However, the underlying species-level OTUs differ between the mesophilic and thermophilic reactors, indicating that organisms even within the same genus can occupy distinct niches in these systems (**Fig. S8**). It is an important observation that for a substantial proportion of the abundant genus-level taxa nothing is known of their potential role in these systems. These include the MiDAS taxa T78, B55_F, and G35_D8, within the phyla Chloroflexi, Firmicutes, and Bacteroidetes, respectively (**Fig. 3B**), which are obvious targets for future research into the ecology of these systems. Influent populations of the archaeal domain were not assessed.

The dominant Archaea in the mesophilic reactors running on primary and surplus sludge was *Methanosaeta*, with a range of other hydrogenoclastic organisms such as *Methanolinea*, *Methanospirillum, Methanobrevibacter* as well as *Ca.* Methanofastidiosa (WCHA1-57) at lower abundances. The uncultured *Ca.* Methanofastidiosa is suggested to be restricted to methylated thiol reduction for methane generation as all known genomes lack genes for acetoclastic and CO_2_-reducing methanogenesis^24^. The dominance of *Methanosaeta* in mesophilic digesters is supported by other studies using amplicon sequencing, qPCR, and shotgun sequencing^7,9,37^.

*Methanothermobacter* and *Methanosarcina* were the dominant acetoclastic methanogens in the thermophilic systems. The difference between the dominant acetoclastic methanogen could be due to process temperature or shorter residence times as both *Methanosaeta* and *Methanosarcina* cover species able to grow across the entire temperature range of operation^38^. Interestingly, *Methanobrevibacter* was also seemingly abundant in the thermophilic reactors, although it is usually considered mesophilic. However, it was not found in mesophilic reactors with thermal hydrolysis pre-treatment, and *Methanobrevibacter* has previously been found in wastewater treatment processes and isolated from faeces^39–41^ – indicating after all that there may be some influence of immigration on archaeal populations in digesters. In addition to a high abundance of *Methanosaeta*, the mesophilic reactors with thermal hydrolysis pre-treatment also had a high abundance of *Methanoculleus*. The methanogen *Methanoculleus* has previously been related to elevated ammonium levels, a relationship that was also supported by the ammonia levels reported for the THP plants in this study (**Table S1**)^9,42,43^.

In this study, we present a comprehensive list of the active microorganisms of full-scale anaerobic digesters receiving primary and surplus sludge from wastewater treatment plants (**Fig. S3**). The relatively low number of genera makes the organisms needed to study feasible, and biological informed decisions less complex and more tractable. Standard application of the curated MiDAS database for anaerobic digester systems, located at wastewater treatment plants, will form an important foundation for future studies of the ecology of these biotechnologically important systems. However, it is important to keep in mind that the list will likely be missing some of the important players due to PCR biases^44^ and that we need primer-free alternatives to get the entire picture of the microbial diversity in anaerobic digesters^45^.

## Materials and methods

### Sampling

Biomass samples from digesters were obtained 2-4 times a year in the period 2011-2016 from 32 ADs at 24 Danish WWTPs (Supplementary Table 1). For primary sludge, 121 samples from 14 WWTPs were sampled during 3 months in October-December, 2014. Each sample was based on a flow proportional collected through 1 day. For surplus activated sludge, 137 sludge samples were obtained from the aeration tank from all 24 WWTPs throughout the 5 years. All samples were homogenised and stored as 2 mL aliquots at -80°C for DNA extraction.

### DNA extraction

DNA was extracted from biomass samples using the FastDNA® Spin kit for soil (MP Biomedicals, Santa Ana, CA, USA) following the standard protocol, except for a 4-time increase in the bead beating duration- as recommended by Albertsen et al., (2015)^21^. The biomass input volume was 50 µl for AD sludge and 500 µl for primary sludge and activated sludge. Primary sludge samples were first filtered onto 0.2-µm pore size polycarbonate filters and the DNA extracted from these using the same method described for other samples.

### DNA amplification and sequencing

#### Bacterial PCR

The bacterial primers used were 27F (AGAGTTTGATCCTGGCTCAG^46^) and 534R (ATTACCGCGGCTGCTGG^47^), which amplify a DNA fragment of ~500 bp of the 16S rRNA gene (variable regions 1-3). 25 µL PCR reactions in duplicate were run for each sample using 1X Platinum® High fidelity buffer, 100 µM of each dNTP, 1.5 mM MgSO_4_, 1 U Platinum® Taq DNA Polymerase High Fidelity (Thermo Fisher Scientific, USA), 400 nM of each barcoded V1-V3 primer, and 10 ng template DNA. PCR conditions were 95°C, for 2 min followed by 30 cycles of {95°C, for 20 s, 56°C for 30 s, 72°C for 60 s} and a final step of elongation at 72°C for 5 min. PCR products were purified using Agencourt AmpureXP (Beckman Coulter, USA) with a ratio of 0.8 bead solution to PCR solution.

#### Archaeal PCR

The archaeal primers used were 340F (CCCTAHGGGGYGCASCA^48^) and 915R (GWGCYCCCCCGYCAATTC^48^), which amplify a DNA fragment of ~ 560 bp of the 16S rRNA gene (variable regions 3-5). 25 µL PCR reactions in duplicate were run for each sample using 1X Platinum® High fidelity buffer, 100 µM of each dNTP, 1.5 mM MgSO_4_, 1 U Platinum® Taq DNA Polymerase High Fidelity (Thermo Fisher Scientific, USA), 400 nM of each V3-V5 primer mix, and 10 ng template DNA. PCR conditions were 95°C, for 2 min followed by 35 cycles of {95°C, for 20 s, 50°C for 30 s, 72°C for 60 s} and a final step of elongation at 72°C for 5 min. PCR products were purified using Agencourt AmpureXP (Beckman Coulter, USA) with a ratio of 0.8 bead solution/PCR solution. Illumina adapters and barcodes were added with a second PCR. 2 µL purified PCR product from above was used as template for a 25 µL PCR reaction containing 1X PCRBIO Reaction buffer, PCRBIO HiFi Polymerase (PCR Biosystems, United Kingdom). PCR conditions were 95°C, for 2 min, 8 cycles of {95°C, for 20 s, 55°C for 30 s, 72°C for 60 s} and a final step of elongation at 72°C for 5 min.

#### Sequencing

Bacteria and archaea amplicon libraries were pooled separately in equimolar concentrations and diluted to 4 nM. The amplicon libraries were paired-end sequenced (2 x 300 bp) on the Illumina MiSeq using v3 chemistry (Illumina, USA). 10-20% PhiX control library was added to mitigate low diversity library effects.

## Read processing and classification

The read data were processed separately for the bacterial and archaeal analysis.

### Bacteria

The paired end reads for the bacterial libraries were trimmed using trimmomatic^49^ and then merged using FLASH^50^. Bacterial reads were screened for potential PhiX contamination using USEARCH (v. v7.0.1090)^51^. The reads were clustered at 97% similarity using USEARCH and subsequently classified using the RDP classifier^52^ with the MiDAS database. The most abundant bacterial (top 80) OTUs from the mesophilic and thermophilic digesters were used to guide curation of the Silva database NR99 v. 1.23 taxonomy as described previously^22^. The resulting updated MiDAS taxonomy (v. 2.1), covering the abundant organisms of both anaerobic digesters and activated sludge, was applied for all analyses presented in this study.

### Archaea

The size of the archaeal V3-V5 fragments made it unattainable to merge the reads, so only read 1 files were used for the analysis. The reads were trimmed to a length of 275 bp. Archaeal reads were screened for potential PhiX contamination using USEARCH (v. v7.0.1090)^51^. The reads were clustered at 97% similarity using USEARCH and subsequently classified using the RDP classifier^52^ with the MiDAS database. The most abundant archaeal OTUs (top 40) from the mesophilic and thermophilic digesters were used to guide curation of the Silva database NR99 v. 1.23 taxonomy as described previously^22^. The resulting updated MiDAS taxonomy (v. 2.1), covering the abundant organisms of both anaerobic digesters and activated sludge, was applied for all analyses presented in this study.

### Data visualisation

Further processing of the OTU table was carried out in the R environment (v. 3.3.2)^53^ using the R studio IDE^54^ using the ampvis package (v. 1.27.0^21^) for visualisation. The ampvis package wraps a number of packages including the phyloseq package (v. 1.19.1)^55^, ggplot2 (v. 2.2.1), reshape2 (v. 1.4.2)^56^, dplyr (v. 0.5.0)^57^, vegan (v. 2.4-1)^58^, knitr (v. 1.15.1)^59^, Biostrings (v. 2.42.1)^60^, data.table (v. 1.10.0)^61^, DESeq2 (v. 1.14.1)^62^, ggdendro (v. 0.1-20)^63^, and stringr (v. 1.1.0)^64^, and cowplot (v. 0.7.0). The samples were subsampled to an even depth of 10 000 reads per sample. Archaeal primers were not specific to the domain, so sequences not classified as Archaea were discarded and the count transformed to a fraction of the archaeal reads. Ratios were calculated between the average abundance for a given OTU within the sample group (mesophilic digesters, thermophilic digesters, mesophilic digesters with thermal hydrolysis pre-treatment) and the average abundance in the influent streams (primary and surplus sludge).

## Data availability

Amplicon sequencing data is available at the ENA with the project ID PRJEB15624. OTU tables and metadata files are available at figshare (DOI: 10.6084/m9.figshare.4308191). The RMarkdown files to generate the figures are available at github (github.com/Kirk3gaard/Publications/tree/master/Kirkegaard2017). The curated MiDAS taxonomy (v. 2.1) is available for download from the MiDAS website (midasfieldguide.org/en/download/).

## Acknowledgements

We would like to express our gratitude for the plant operators for sending samples and supplying metadata. The project was funded by the Villum foundation (grant no. VKR 022796) and the Innovation Fund Denmark (NomiGas, grant no. 1305-00018B) to P.H. Nielsen. S. McIlroy was supported by the Danish Council for Independent Research (grant no. 4093-00127A).

## Supplementary

**Figure S1.**
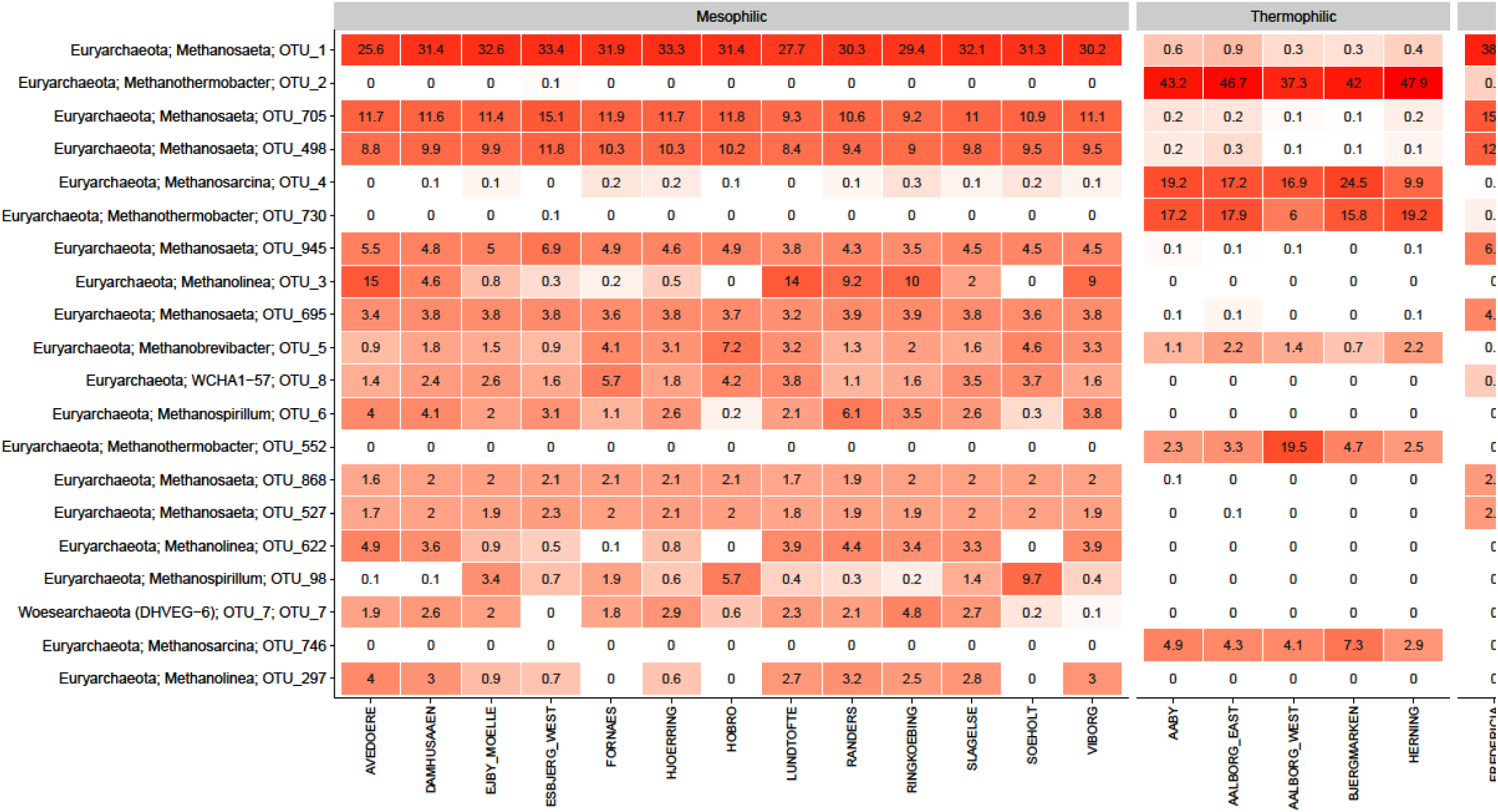
Heatmap of the 20 most abundant archaeal OTUs. Phylum and genus level classifications of the OTUs are shown, when no classification is available OTU number is given. The wastewater treatment plants had a total of 32 AD reactors (1-4 per plant) and they were analysed 2-23 times. The mean abundance is shown for each plant. The taxa are sorted by mean abundance across the plants.

**Figure S2.**
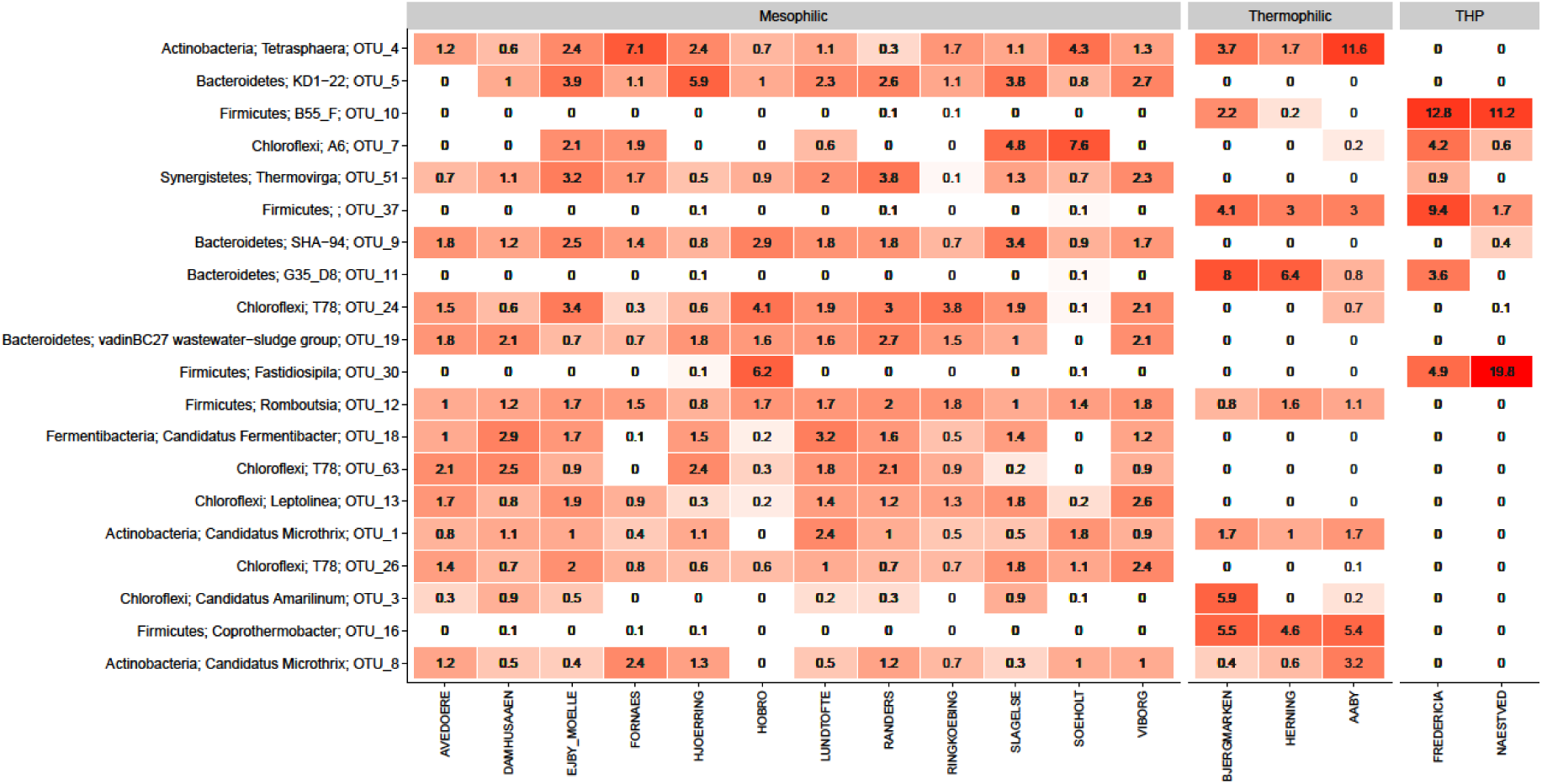
Heatmap of the 20 most abundant bacterial OTUs. Phylum and genus level classifications of the OTUs are shown, when no classification is available the field is empty. The wastewater treatment plants had a total of 32 AD reactors (1-4 per plant) and they were analysed 3-30 times. The mean abundance is shown for each plant. The taxa are sorted by mean abundance across the plants.

**Figure S3.**
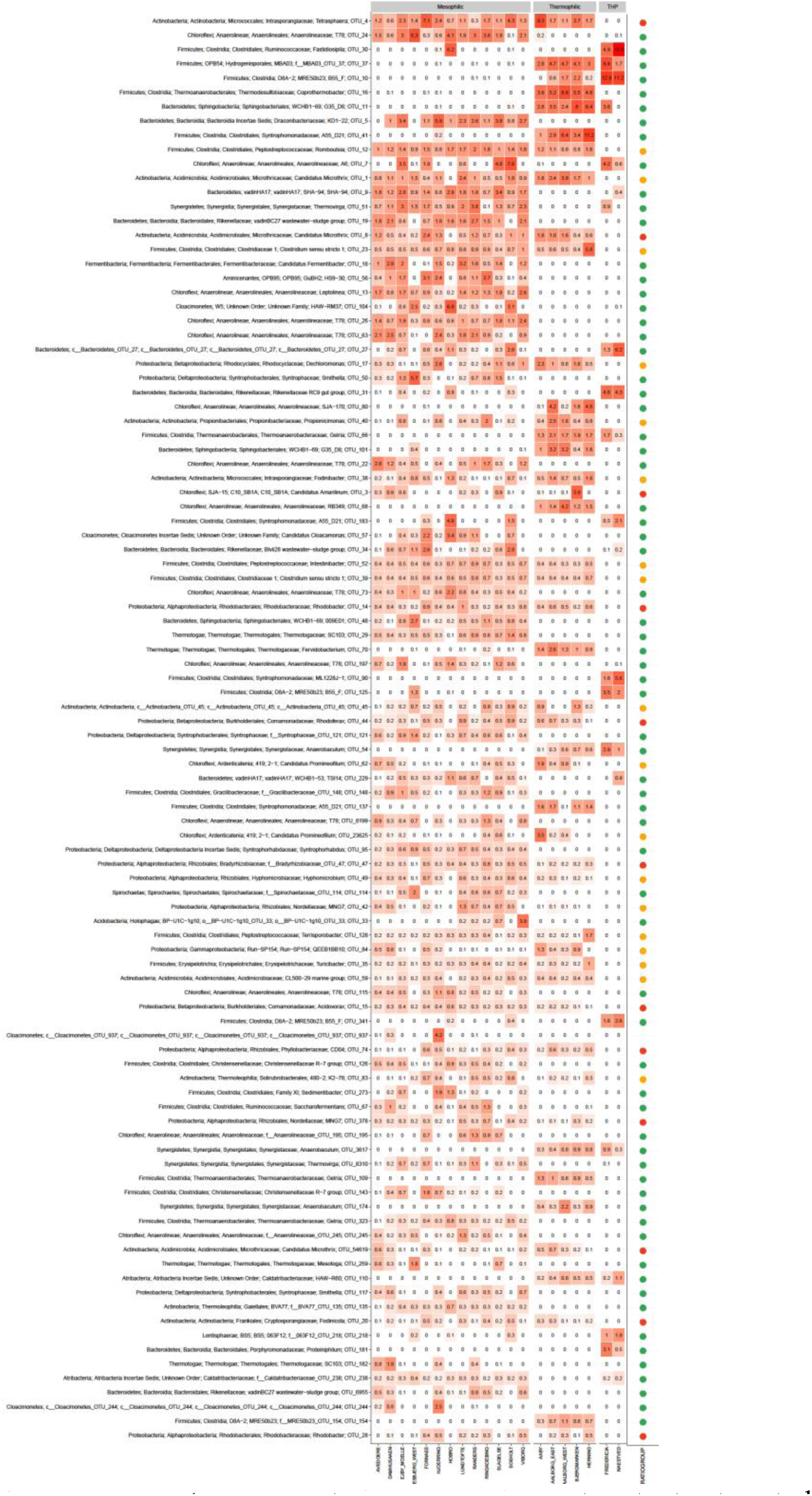
Heatmap of the 100 most abundant bacterial OTUs. The wastewater treatment plants had a total of 32 AD reactors (1-4 per plant) and they were analysed 3-30 times. The mean abundance is shown for each plant. The taxa are sorted by mean abundance across the plants. The right panel indicates whether the digester to influent abundance ratio was above 10 (•) and likely actively growing, between 1 and 10 (•), or below 1 (•), indicating no growth.

**Figure S4.**
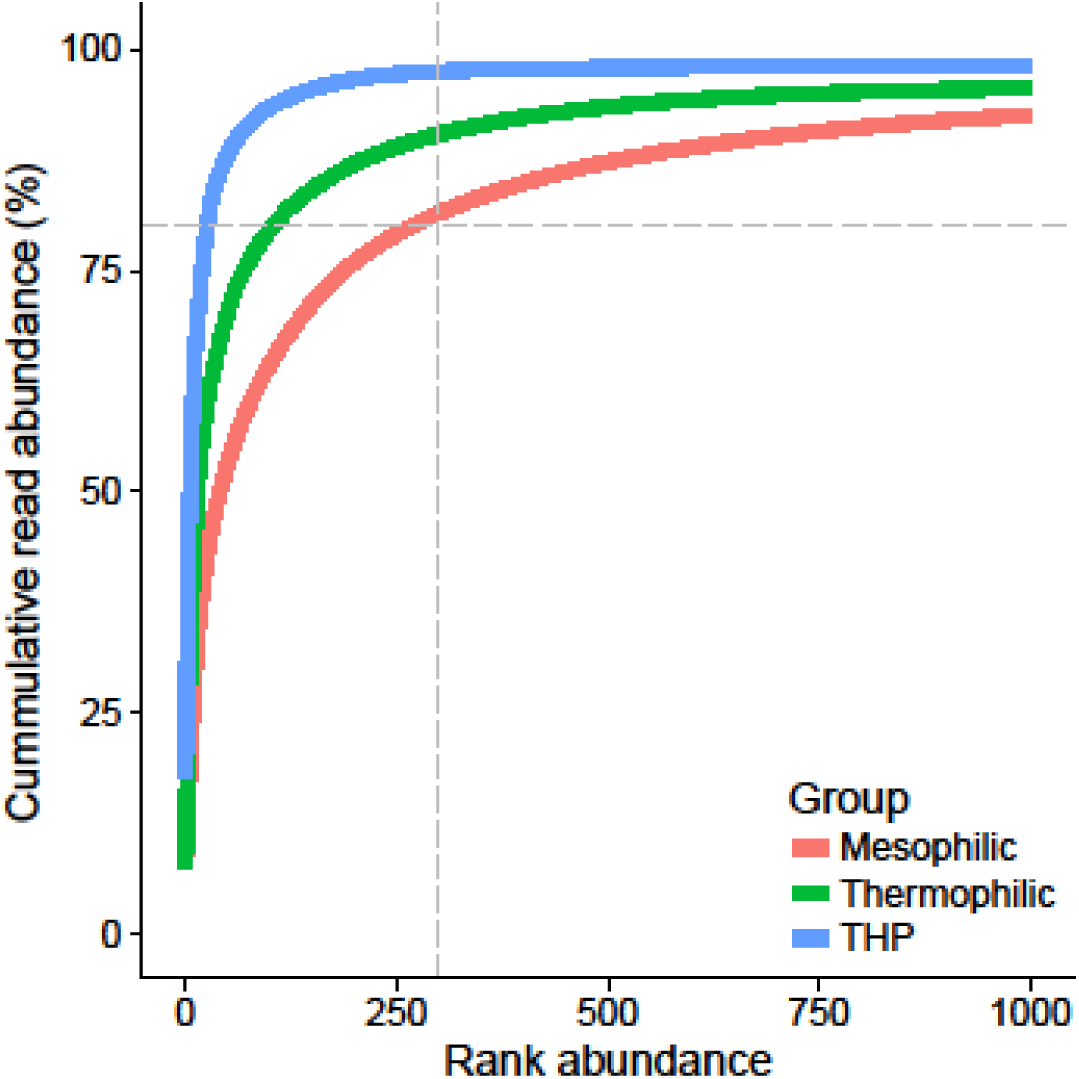
Rank abundance curve indicating the number of OTUs needed to account for a certain fraction of the cumulative reads for mesophilic (■),thermophilic (■) and mesophilic with THP (■) samples.

**Figure S5.**
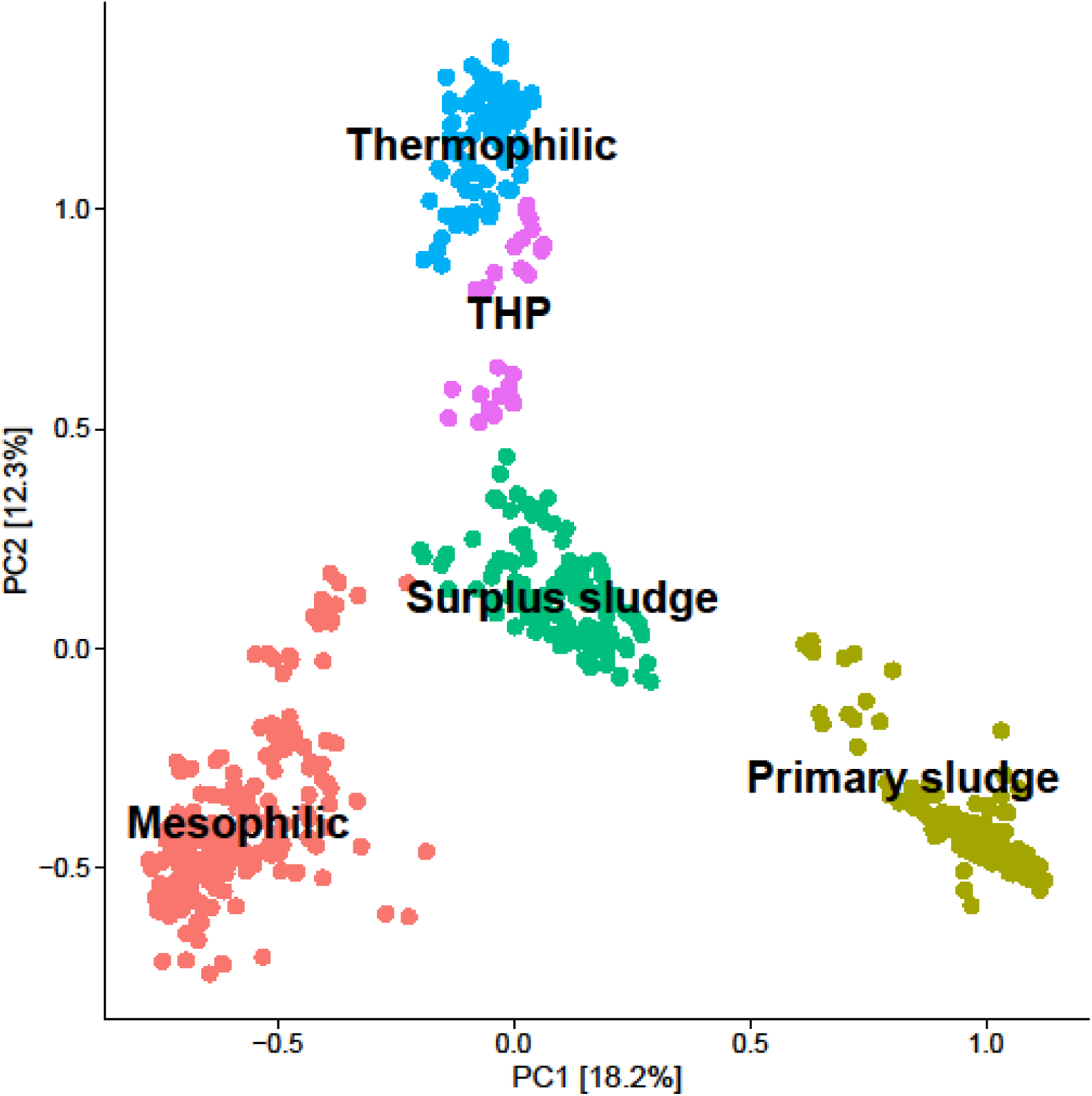
Principal component analysis of the bacterial communities analysed in this study highlighting samples by process type information. Mesophilic (•), thermophilic (•), and mesophilic with thermal hydrolysis pre-treatment (• THP), primary sludge (•), and surplus sludge (•).

**Figure S6.**
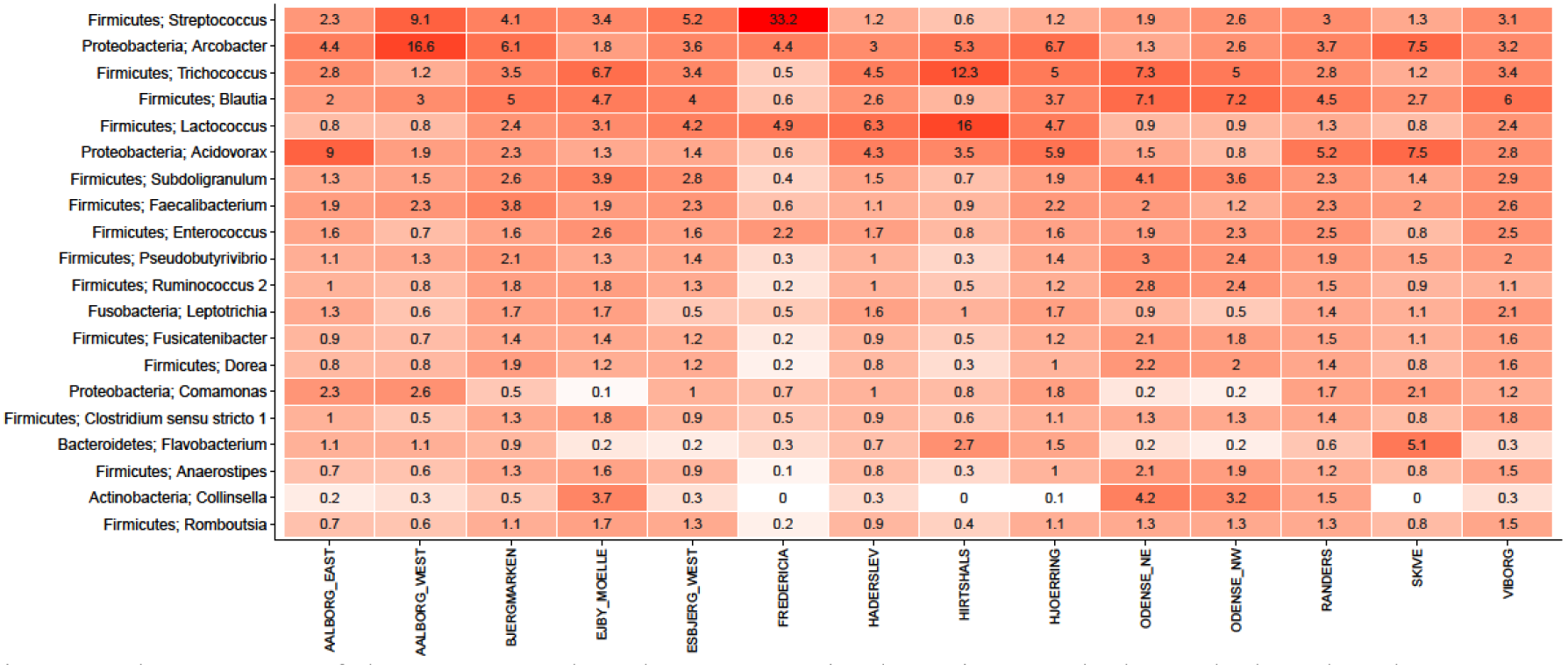
Heatmap of the 20 most abundant genera in the primary sludge. Phylum level classifications are shown. The mean abundances are shown for each plant (3-23 samples per plant). The taxa are sorted by mean abundance across the plants.

**Figure S7.**
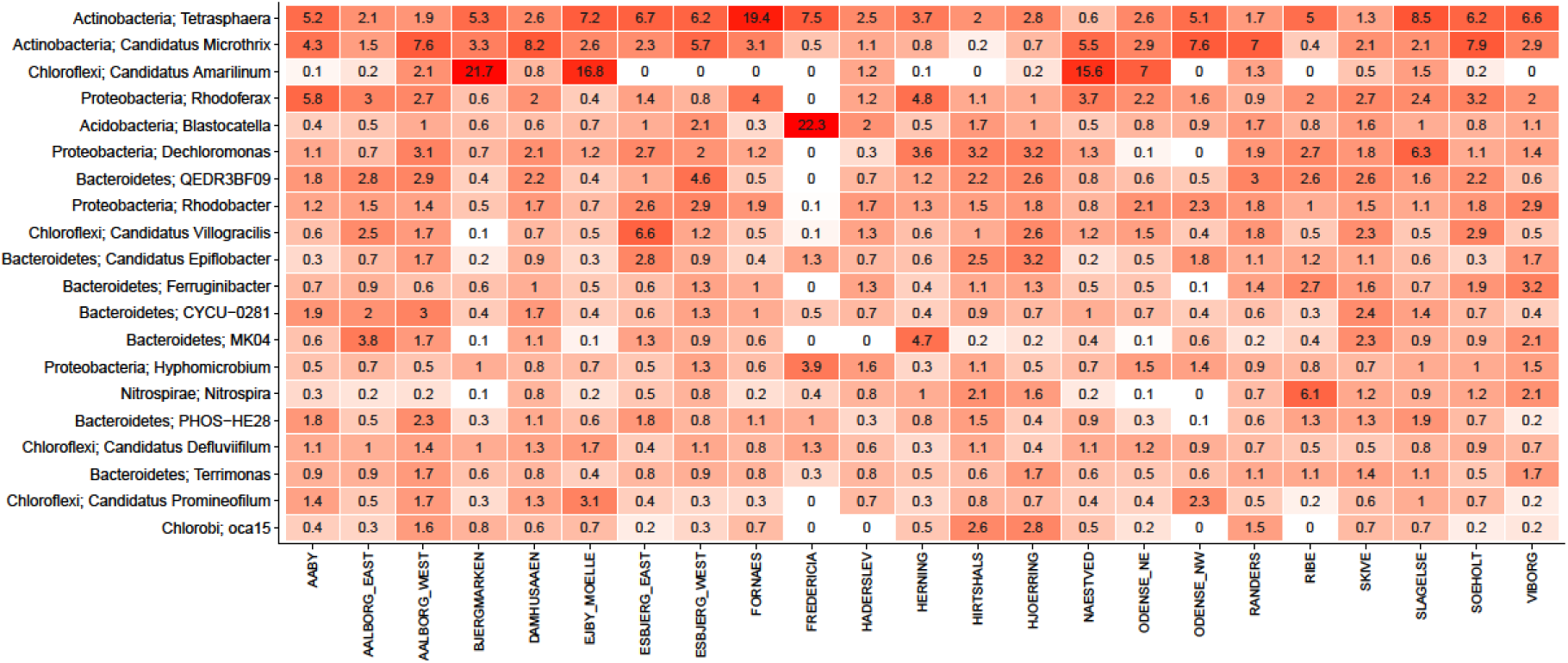
Heatmap of the 20 most abundant genera in the surplus sludge. Phylum level classifications are shown. The mean abundance is shown for plants with more than 1 sample (1-24 samples per plant). The taxa are sorted by mean abundance across the plants.

**Figure S8.**
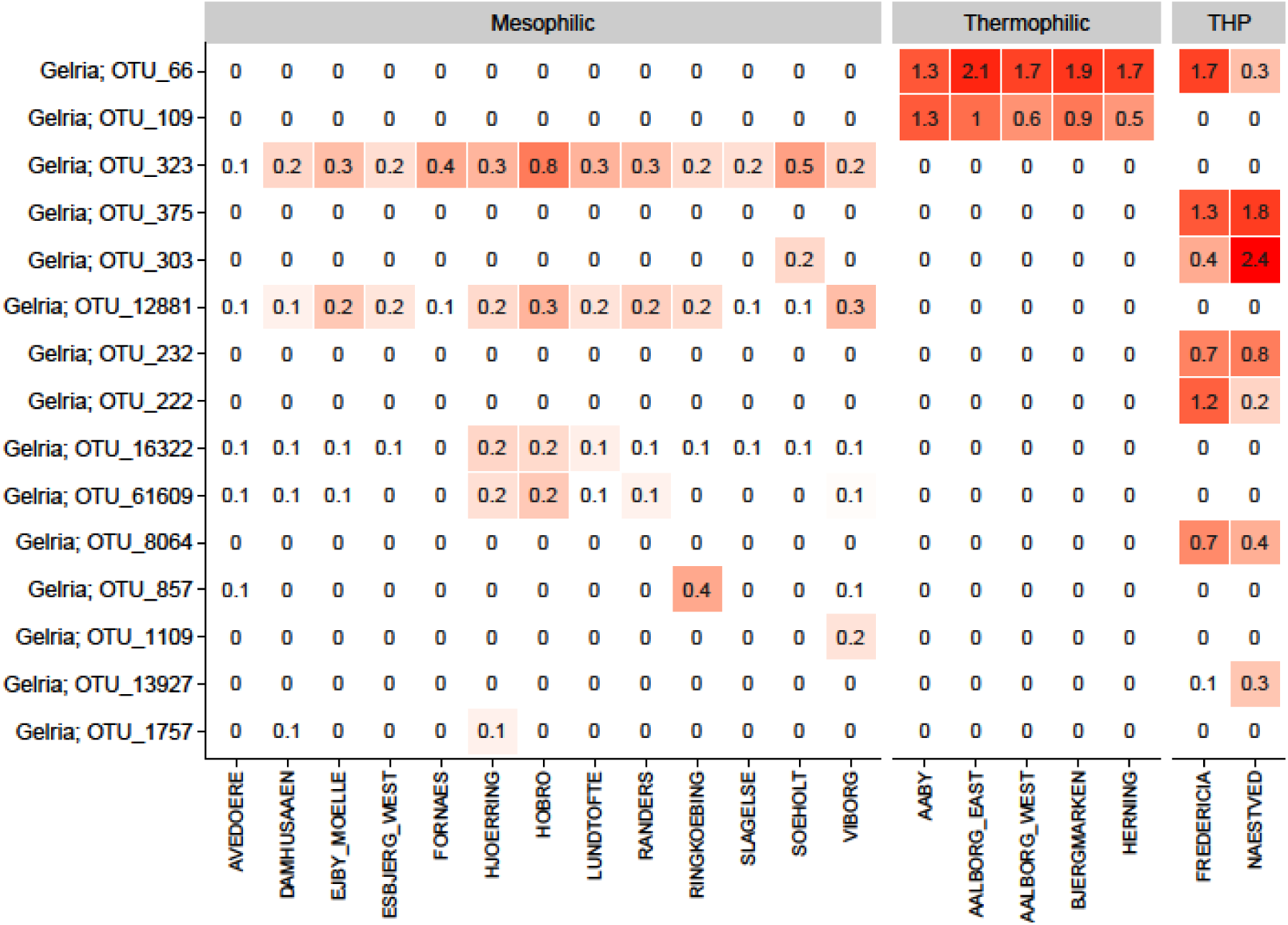
Heatmap of the 15 most abundant OTUs belonging to the genus *Gelria* in mesophilic and thermophilic systems. The wastewater treatment plants had a total of 32 AD reactors (1-4 per plant) and they were analysed 3-30 times. The mean abundance is shown for each plant. The taxa are sorted by mean abundance across the plants.

### Table S1

Table S1 | Plant overview: locations, process types, temperatures etc. Moved as it is not suitable for word

### Table S2

Table S2 | Digester to influent ratios, and mean abundance values at the OTU level for mesophilic, thermophilic and mesophilic with THP. Moved as it is not suitable for word

